# Targeted brain-specific tauopathy compromises peripheral skeletal muscle integrity and function

**DOI:** 10.1101/2023.11.17.567586

**Authors:** Bryan Alava, Gabriela Hery, Silvana Sidhom, Stefan Prokop, Karyn Esser, Jose Abisambra

**Affiliations:** Department of Physiology and Aging, University of Florida, Gainesville, Florida, 32610, USA; Center for Translational Research in Neurodegenerative Disease (CTRND), University of Florida, Gainesville, Florida, 32610, USA; Department of Neuroscience, University of Florida, Gainesville, Florida, 32610, USA; Department of Pathology, University of Florida, Gainesville, Florida, 32610, USA; Brain Injury Rehabilitation and Neuroresilience (BRAIN) Center, University of Florida, Gainesville, Florida, 32601, USA

## Abstract

Tauopathies are neurodegenerative disorders in which the pathological intracellular aggregation of the protein tau causes cognitive deficits. Additionally, clinical studies report muscle weakness in populations with tauopathy. However, whether neuronal pathological tau species confer muscle weakness, and whether skeletal muscle maintains contractile capacity in primary tauopathy remains unknown. Here, we identified skeletal muscle abnormalities in a mouse model of primary tauopathy, expressing human mutant P301L-tau using adeno-associated virus serotype 8 (AAV8). AAV8-P301L mice showed grip strength deficits, hyperactivity, and abnormal histological features of skeletal muscle. Additionally, spatially resolved gene expression of muscle cross sections were altered in AAV8-P301L myofibers. Transcriptional changes showed alterations of genes encoding sarcomeric proteins, proposing a weakness phenotype. Strikingly, specific force of the soleus muscle was blunted in AAV8-P301L tau male mice. Our findings suggest tauopathy has peripheral consequences in skeletal muscle that contribute to weakness in tauopathy.

## Introduction

Tauopathies, including Alzheimer’s disease (AD) and frontotemporal dementias associated with chromosome 17 (FTD), are more than 20 neurodegenerative disorders that pathologically feature intracellular aggregates of the protein tau termed neurofibrillary tangles (NFTs) ^1–5^. Tau canonically binds and stabilizes microtubules ^5–7^; however, pathogenic processes disrupt tau-microtubule complexes. Emerging data characterizing tau interactomes suggest that tau performs functions beyond stabilizing microtubules, and therefore, alternative mechanisms may mediate tau pathogenic processes ^8–16^

Besides cognitive decline, muscle weakness also manifests early in tauopathies ^17–19^. Moreover, murine models exhibiting AD-related amyloid pathology in the brain or targeted to skeletal muscle ^20,21^ show changes in skeletal muscle features such as specific force, fiber size, and muscle gene expression. These data link muscle weakness with AD, a tauopathy with amyloid plaque co-pathology, but it is not yet understood to what extent primary tau pathology independently impacts skeletal muscle. To explore the association between muscle weakness and onset of tau pathology, we investigated whether pathological tau species in the brain link to skeletal muscle abnormalities in a model of primary tauopathy.

Murine models, such as the JNPL3, PS19, and rTg4510 ^22–27^ models manifest motor deficits and muscle anomalies. However, whether these peripheral abnormalities are the consequence of brain-specific overexpression of pathological tau species, and whether muscle is impacted by tauopathy independent of peripheral nerve function, remains unknown. In this study, we performed an extensive evaluation of muscle structure and function in AAV8-P301L tau expressing mice. We found skeletal muscle abnormalities with the potential to induce physiological consequences, implicating skeletal muscle pathology and dysfunction as a peripheral symptom of tauopathy.

## Results

### Intraventricular AAV8-P301L tau expression promotes widespread pathology

We first validated that the PND0 intracerebroventricular brain injections of P301L tau induced tauopathy by harvesting 15-month-old AAV8-P301L tau-injected mouse brains (Fig. 1). We identified widespread and consistent human tau expression, which aligned with previously published data using ICV injections of AAV1-P301L tau ^1^. Distribution was largely concentrated in the hippocampus, cortex, and olfactory bulb (Fig. 1), as expected. Limited immunoreactivity was observed in the cerebellum, suggesting fine motor control directed by the cerebellum may not be directly impacted by AAV8-guided P301L tau expression in this model. We also assessed accumulation of insoluble tau using biochemical analyses of soluble and sarkosyl-insoluble cortical tau. We identified increased immunoreactivity to pathology-associated pS202/T205 tau (AT8) in cortical samples of AAV8-P301L mice (Supplemental Fig. 1A, E, H). As expected, total tau (endogenous murine and AAV8-driven human) was also increased in AAV8-P301L mice (Supplemental Fig. 1B and 1G). We identified accumulation of soluble (Supplemental Fig. 1C, F) and sarkosyl-insoluble human-derived tau in cortical samples from the P3 pellet fraction (Supplemental Fig. 1D). These data provide histological and biochemical evidence that AAV8-P301L tau injected mice present neuropathological brain features of tauopathy.

**Fig. 1.**
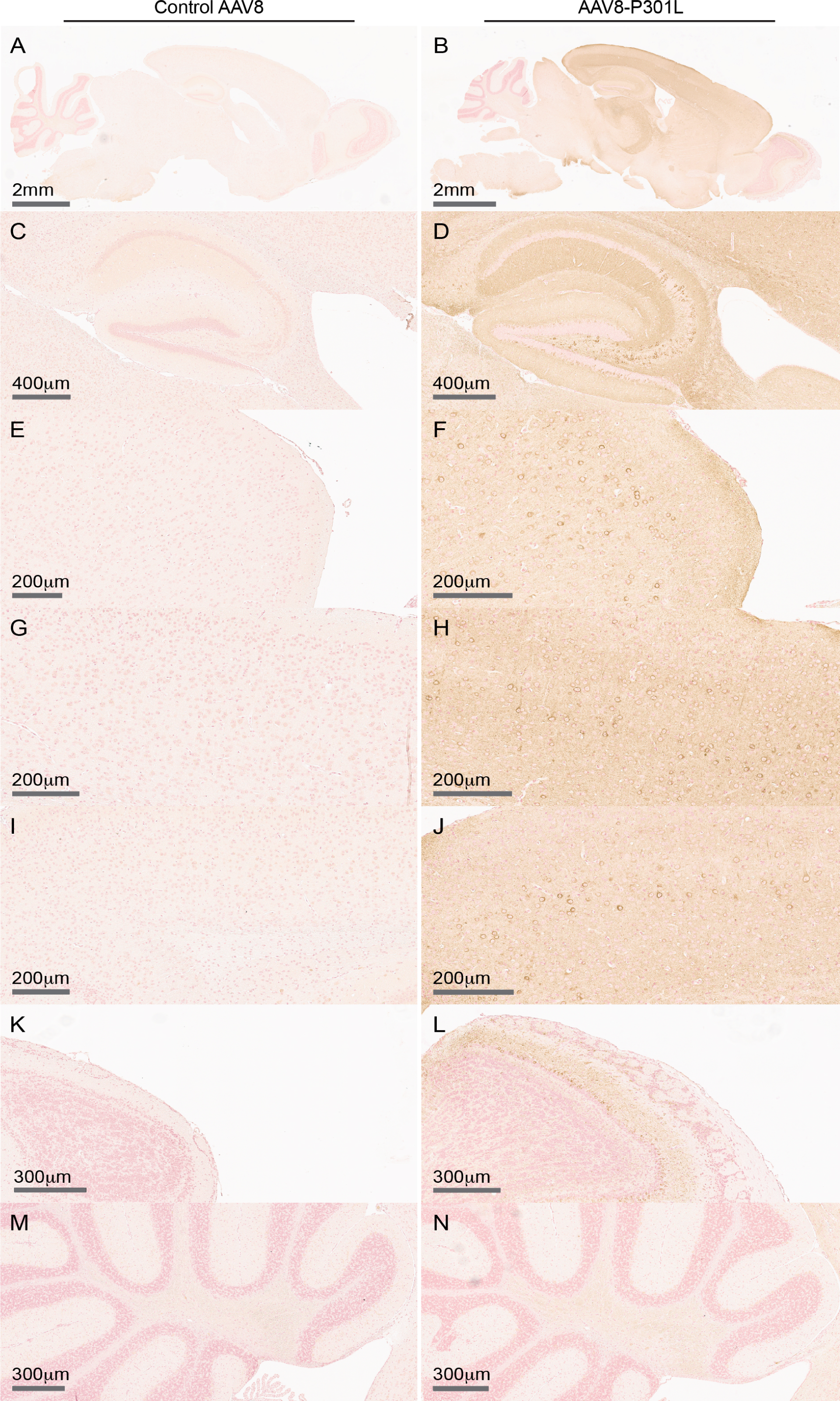
AAV8-P301L tau expression develops widespread tangle pathology. Compared to AAV8 controls (A,C,E,G,I,K,M), P301L-tau was detected in AAV8-P301L tau mouse brains (B). Human tau was largely detected in AAV8-P301L tau-expressing hippocampal, anterior, middle, and posterior cortical regions (D,F,H,J), and the olfactory bulb (F). The cerebellum showed limited P301L-tau immunoreactivity (G). N=4 female, 7 male AAV8-P301L mice, N=5 female, 6 male control AAV8 mice. See also Figure S1.

### AAV8-P301L tau expression promotes abnormal locomotor phenotypes and muscle histological features independent of changes in fiber size

AAV8-P301L tau mice showed increased daily cage activity compared to sex-matched controls at 8 months of age (Fig. 2a) as measured by infrared (IR) cage sensors. This increase is largely driven by activity during the dark/active phase; interestingly, male AAV8-P301L tau mice increased cage activity in both the light/inactive and dark/active phases (Fig. 2B-C). Activity in P301L tau mice remained consistently increased each day during the 8-day recording period (Supplemental Figure 2A-C), yet the timing of activity onset was not altered (Fig. 2D-E).

**Fig. 2.**
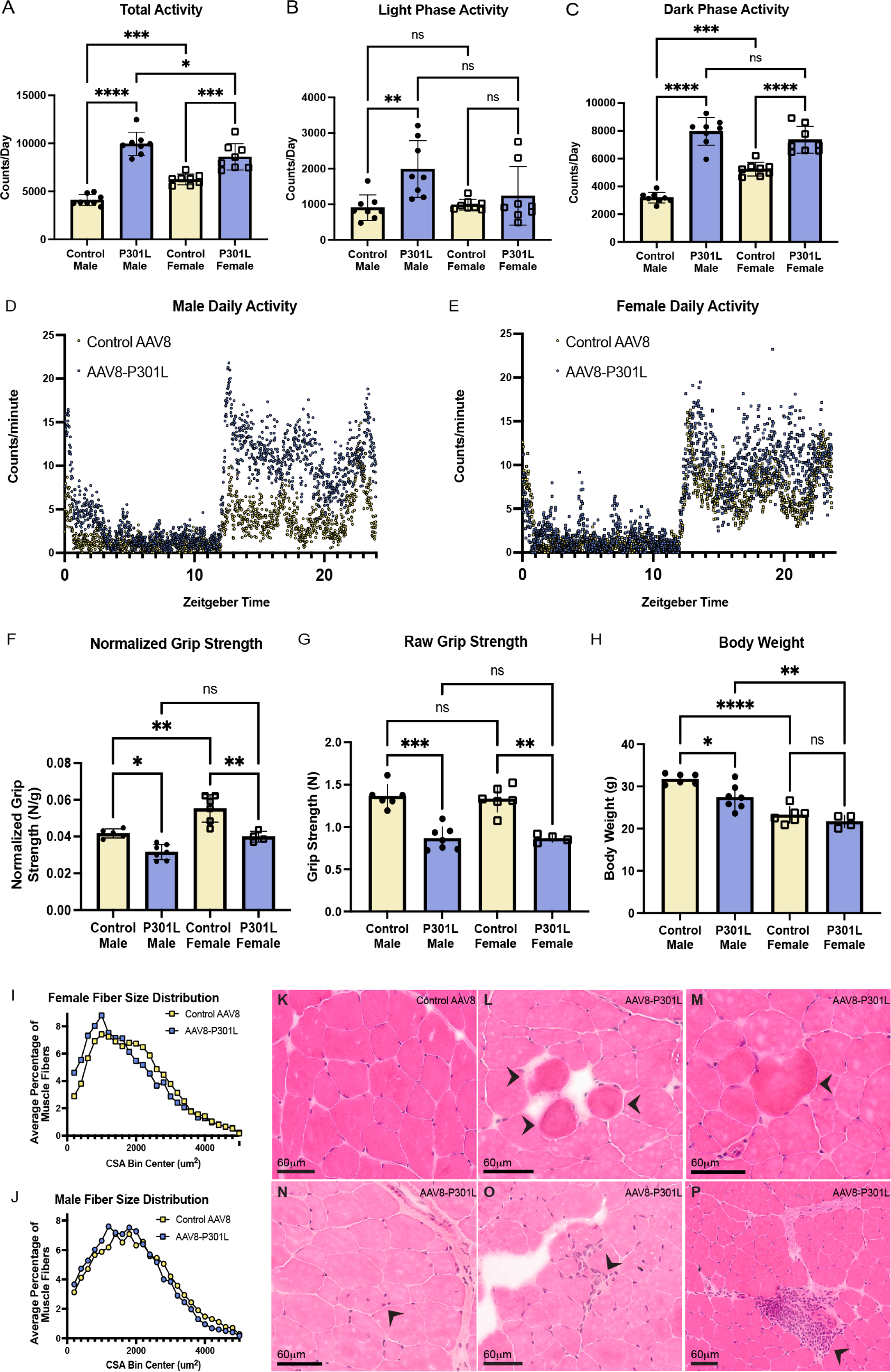
Abnormal locomotor phenotypes and histological features persist in AAV8-P301L tau mice independent of changes in fiber size. A-E) AAV8-P301L tau mice are more active (A). Only male AAV8-P301L tau mice are more active in the light/rest phase compared to controls (B). Both male and female AAV8-P301L tau mice are more active in the dark/active phase compared to their sex matched controls (C). Averaged counts per minute for 7 days show increases in activity and no changes in timing of activity (D, E). N=4 female, 7 male AAV8-P301L mice, N=5 female, 6 male control AAV8 mice; Ordinary two-way ANOVA, \**p* < 0.05, ** *p* < 0.01, *** *p* < 0.001, *** *p*< 0.0001. See also Figure S2. F-H) Grip strength normalized to body weight is significantly reduced in AAV8-P301L male and female mice compared to their respective controls (A). Raw grip strength is reduced in the AAV8-P301L tau mice (B). Body weight was only significantly reduced in P301L tau expressing males (C). N=4 female, 7 male AAV8-P301L mice, N=5 female, 6 male control AAV8 mice; Ordinary two-way ANOVA, * *p* < 0.05, ** *p* < 0.01, *** *p* < 0.001. I-J) Calculated cross-sectional area distribution is unchanged in AAV8-P301L mice compared their sex-matched controls (A, B). N=4 female, 5 male AAV8-P301L mice, N=6 female, 6 male control AAV8 mice. See also Figure S2. K-P) Compared to wildtype mice (A), AAV8-P301L tau mice displayed subtle abnormalities in fiber shape and structure (circular fibers (B,C), group of smaller myofibers with central nuclei (D)) and abnormal clusters of nuclei ranging in severity (potentially pathogenic (E), and noticeably severe (F)).

To determine changes in skeletal muscle due to tauopathy, grip strength was measured at 3 months of age, which is 3 months prior to the appearance of cognitive deficits previously established in this model ^1^. P301L tau mice of both sexes showed grip strength deficits (Fig. 1F, G). The deficit in grip strength normalized to body weight (Fig. 2F) is driven largely by a reduction in raw grip strength, which was reduced in AAV8-P301L tau mice of both sexes (Fig. 2G). Surprisingly, body weight was significantly reduced in AAV8-P301L tau-expressing males but not females (Fig. 2H). These data suggest that this model manifests muscle weakness independently of their body weight.

To assess if general weakness was driven by changes in muscle size, we measured fiber size distribution of the tibialis anterior muscle. We did not observe differences in myofiber calculated cross-sectional area distribution (Fig. 2I, J) or minimum feret diameter distribution (Supplemental Fig. 2D, E). Therefore, altered fiber size distribution does not explain the grip strength weakness observed in this model.

We then assessed gross histological morphology of the tibialis anterior fibers. As expected, muscle sections from AAV8-P301L mice did not show visible signs of myofiber atrophy. Despite minor artifacts from freeze damage, there were common abnormalities in the tibialis anterior muscle of AAV8-P301L tau expressing mice not detected in the control mouse muscles. AAV8-P301L tau mice showed histological signs of myopathy, including circular fibers that deviated from normal polygonal shape (Fig. 2L, M) and relatively small fibers with central nuclei (Fig. 2N). AAV8-P301L tau muscle cross section displayed a preponderant number of central nuclei compared to controls. Darker, circular fibers were detected in 5 of the 8 mice, but not apparent in any controls. The most overt histological feature was the abnormal clustering of nuclei. The abnormal clustering ranged in severity in the cross sections between AAV8-P301L tau expressing mice. The majority of the AAV8-P301L tau mice displayed no nuclear clustering (4 out of the 8 AAV8-P301L mice), consistent with normal muscle (Fig. 2K). However, nuclear clustering was seen in 3 out of the 8 AAV8-P301L tau muscles analyzed (Fig. 2O), and severe nuclei grouping was observed in 1 AAV8-P301L tau muscle cross section (Fig. 2P). The severe nuclei grouping was apparent in the three serial cross sections analyzed. Since cross sections are representative of only the sagittal view of the fibers, there may be additional pathogenic features in the muscle of other mice not captured in the cross sections analyzed. Therefore, distinctions in AAV8-P301L myofibers observed with H&E analysis was strengthened with a spatial transcriptomic study.

### Altered transcript detection in the tibialis anterior of AAV8-P301L tau expressing mice is sex and region specific

To determine if selected regions of myofibers from AAV8-P301L tau-expressing mice were distinct from controls, spatial transcriptomics was performed on cross sections of the tibialis anterior muscle of AAV8-P301L tau mice and AAV8 controls. To compare the localized gene expression patterns, we selected regions of standard myofibers, myofibers with central nuclei, myofibers proximal to vasculature, and myofibers proximal to nerve bundles (Fig. 3A-D). To visualize the regions, we stained the muscle cross sections with laminin and SYTO-13 (Supplementary Figure 3) and selected regions that were in proximity to the section landmarks noted above.

**Fig. 3.**
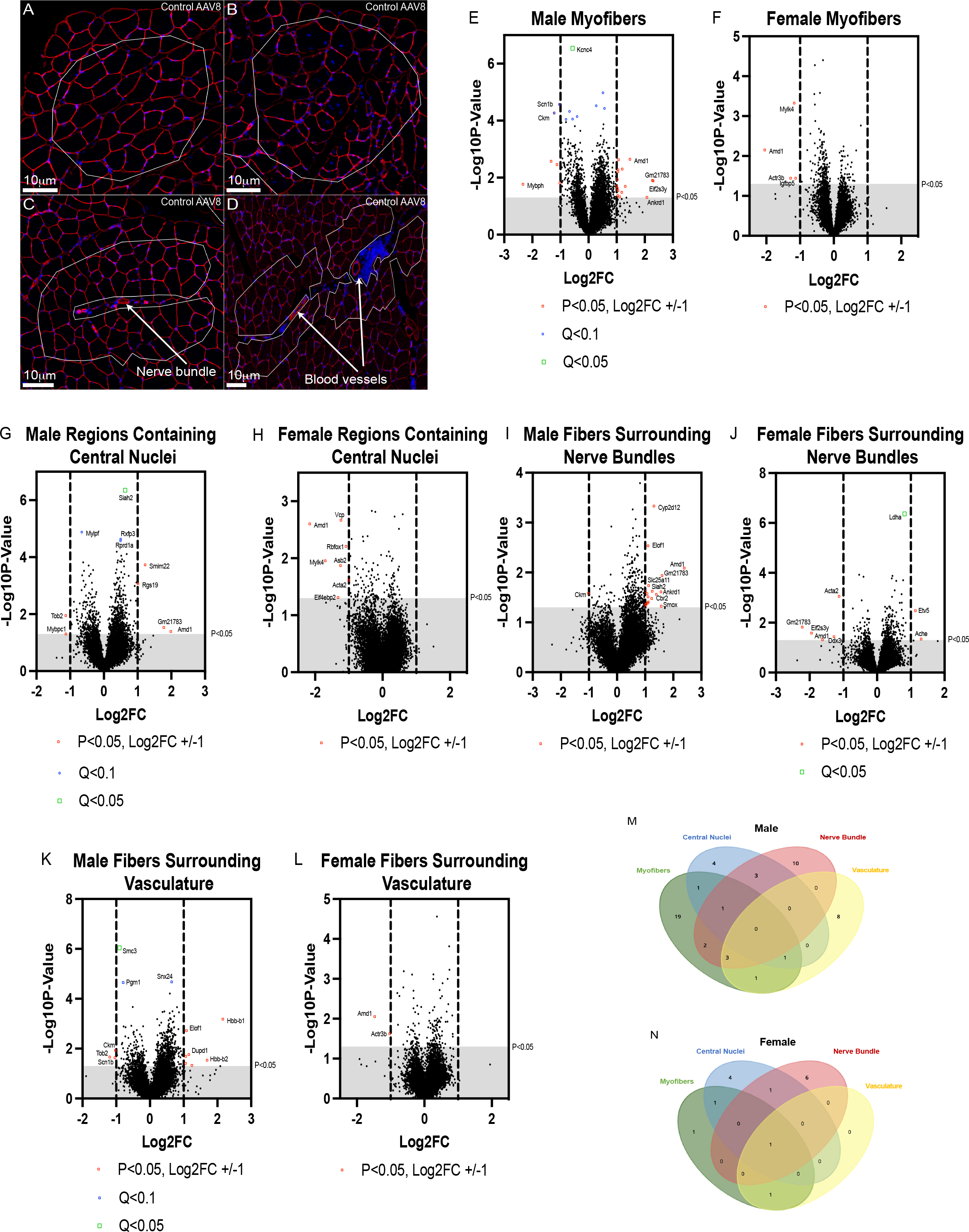
Transcriptional differences in the tibialis anterior cross sections of AAV8-P301L tau mice are sex and muscle hallmark proximity dependent. Representative images of standard myofiber regions (without central nuclei and not directly proximal to a nerve bundle or vasculature) (A), regions of myofibers with central nuclei (B), proximal to nerve bundles (C) proximal to blood vessels (D showing two regions). Differentially detected transcripts for each sex and region type annotated by P_adj_ < 0.1 or P_unadj_ < 0.05 and Log2FC +/-1 (E-L). Region-type and sex specificity of differentially detected transcripts (M, N). Linear Mixed Model with Benjamini-Hochberg correction for P_adj_ values. N = 3 mice/sex/treatment. 2 replicates per region type per mouse. Red: laminin stain; Blue: SYTO-13 stain. See also Figure S3.

We detected altered gene expression in standard myofiber regions in AAV8-P301L mice (Fig. 3 and Supplemental Table 1). Twenty-eight differentially expressed genes (DEGs) were identified in AAV8-P301L males and 4 in AAV8-P301L females (Fig. 3E, F). *Ckm* (Creatine Kinase, an important metabolic enzyme that supports high energy levels of phosphocreatine in skeletal muscle) was surprisingly downregulated in AAV8-P301L males. DEGs encoding sarcomeric proteins such as *Mybph* (Myosin Binding Protein H, downregulated), *Myh4* (Myosin Heavy Chain 2B/fast, downregulated), and *Ankrd1* (Ankyrin Repeat Domain-Containing Protein 1, upregulated) were identified in males, and *Mylk4* (Myosin Light Chain Kinase Family Member 4) was downregulated in females. Beyond sarcomeric proteins, the transcript for calcium sequestering protein 2 (*Casq2 cardiac/slow muscle)* was also upregulated in males. This would increase the calcium buffering capacity in the muscle of the AAV8-P301L muscle. Moreover, the *Scn1b* (Sodium Voltage-Gated Channel Beta Subunit 1) transcript was downregulated in AAV8-P301L males. This encodes a regulatory subunit of the voltage-gated sodium channel, which is essential for action potential initiation and propagation.

In regions containing central nuclei, we found 10 DEGs in male AAV8-P301L tau mice and 7 in females (Fig 3G, H). DEGs encoding sarcomeric proteins were also altered in this region. *Mybpc* (Myosin Binding Protein-C, downregulated) and *Mylpf* (myosin light chain, phosphorylatable, also known as *Mlc2f,* downregulated) were changed in males. *Mylk4* was downregulated in females, paralleling the standard myofiber regions.

We identified the same sex-specific distinctions (increased DEGs in males) reflected in myofibers proximal to blood vessels (Fig. 3I, J). Thirteen DEGs were present in AAV8-P301L males, and only 2 in females. We observed downregulation of *Ckm* and *Scn1b* in males, resembling the standard myofibers.

Increased DEGs were maintained in AAV8-P301L males compared to females in regions of myofibers surrounding nerve bundles (19 vs 8 DEGs) (Fig. 3K, L). Reduced *Ckm* was evident in AAV8-P301L males. The detection of the *Ankrd1* transcript was upregulated in males, reflecting standard myofibers. Interestingly, *Ache* (Acetylcholinesterase) was upregulated in female regions surrounding nerve bundles. Acetylcholinesterase functions in the rapid degradation of synaptic acetylcholine at the neuromuscular junction. Therefore, its upregulation would be consistent with more rapid clearing of acetylcholine at the neuromuscular junction.

Interestingly, *Amd1* (adenosylmethionine decarboxylase 1, polyamine biosynthesis) was differentially detected in 7 of the 8 region types analyzed. *Amd1* was consistently downregulated in females, and consistently upregulated males. Sex specificity to polyamine biosynthesis has been described and may explain this phenomenon ^28^. Most differentially detected transcripts had regional specificity that was determined by sex (Figs. 3M, N). However, many differentially detected transcripts encoded proteins present at the sarcomere or essential in generating force, and they were detected in every region for both sexes.

### Reduction of specific force in the muscle of AAV8-P301L tau male mice

We hypothesized that changes in the levels of transcripts encoding sarcomeric proteins alter force generation in AAV8-P301L tau expressing mice, and that these changes were independent of neuronal signaling. To determine whether limited contractile force production contributes to weakness in AAV8-P301L tau mice, we obtained *ex vivo* measures of maximum isometric tetanic force from two different muscles in the hindlimb: the soleus muscle and the extensor digitorum longus (EDL) muscle. Given that changes in activity and DEG were more pronounced in male mice, we focused primarily in males. *Ex vivo* muscle mechanical measures activate the muscle using field stimulation and reflect the intrinsic force generating capacity of the muscle, which is not indicative of nerve-muscle communication. Therefore, the data acquired in this technique reflects changes in muscle phenotype independently from nerve stimulation. Maximum tetanic force of the soleus was not significantly different between control and AAV8-P301L tau males. Specific force, which is maximum isometric tetanic force normalized for muscle size (physiological cross-sectional area) was significantly reduced in AAV8-P301L tau males compared to controls (Fig. 4A). This finding demonstrates that intrinsic force generating properties of the soleus, an important postural muscle, are impaired in this tauopathy model. Analysis of the soleus force-frequency properties did not detect shifts between the control and AAV8-P301L mice, suggesting that the reduction in specific force is not likely due to any problems with calcium handling/excitation contraction coupling (Fig. 4B).

**Fig. 4.**
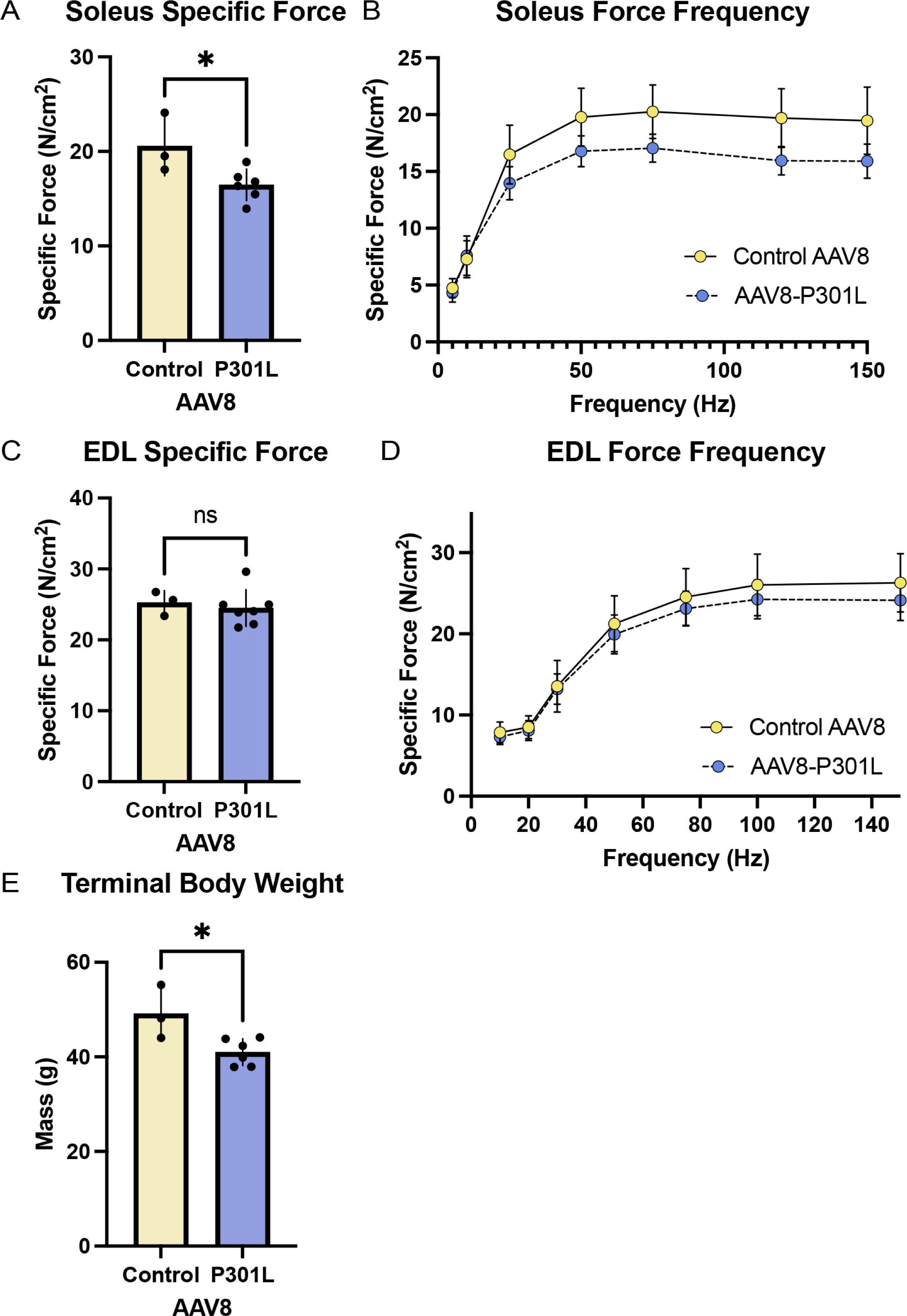
Ex vivo specific force of the soleus is reduced in AAV8-P301L tau expressing males. Soleus *ex vivo* specific force calculations are reduced in the AAV8-P301L tau expressing males (A), and normal force frequency curves are observed (B). Specific force of the EDL muscle is sustained (C-D). Reduced body weight was maintained in AAV8-P301L male mice at time of collection (E). N=6-7 male AAV8-P301L tau mice, N=3 control AAV8 male mice; Student’s t-test, *P< 0.05. See also Figure S4.

We also determined specific force for the EDL from AAV8-P301L tau males, which was not significantly different from control males (Fig. 4C). Reflecting the soleus, the force frequency curves were similar, indicating normal calcium handling (Fig. 4D). Reduced body weight was maintained at 15 months in AAV8-P301L males (Fig. 4E).

Since the tauopathy model is generated with an exogenous ICV injection of AAV8-P301L tau, the decrease in specific force of the soleus muscle could have been due to an accumulation of human tau in the soleus. This could sterically impact normal sarcomeric structure or protein-protein interactions (i.e., actin-myosin), and limit myofiber force production. However, detectable accumulation of human tau was not observed in the soleus of AAV8-P301L tau expressing mice (Supplemental Figure 4). Therefore, an alternative mechanism may be responsible for specific force deficits in AAV8-P301L tau expressing males.

## Discussion

We identified skeletal muscle abnormalities in the AAV8-P301L tauopathy mouse model as an addition to the growing evidence of skeletal muscle irregularities as peripheral symptoms of tauopathy ^20,21,27^. This work is the first to emphasize skeletal muscle anomalies due to brain-specific overexpression of P301L tau inducing tauopathy.

The AAV8-P301L tau model is ideal to study muscle weakness in tauopathy because P301L tau is introduced via ICV injection under the human synapsin 1 promoter packaged in recombinant AAV8, and therefore is not endogenously overexpressed in skeletal muscle fibers (Supplemental Fig. 4). Overexpression of mutant tau in skeletal muscle could be an unintentional consequence of tauopathy models under the mouse prion promoter ^23,24,29^. In the AAV8-P301L model, *in vivo* weakness is observed prior to reported cognitive deficits of the model, which aligns with the clinical increased risk for dementia in humans with low handgrip strength^17,30–33^. Furthermore, human tau overexpression is not detected in muscle tissue and *ex vivo* muscle contractile weakness was detected independent of motor neuron signaling (Fig. 4 and Supplemental Fig. 4). These findings warrant additional studies that can be leveraged using the AAV8-P301L model.

The deficit on the grip strength meter observed in AAV8-P301L tau-expressing mice merited investigation into skeletal muscle properties that may underpin this finding. We observed abnormalities such as mononuclear cell infiltration in H&E-stained muscle cross sections that could have been specific to the cross-section or throughout the length of the muscle. Therefore, we performed spatial transcriptomic sequencing to identify the nature of the cellular proliferative process in the fibers themselves that was not apparent in the H&E-stained sections. We observed sex specific transcriptional changes in the myofibers of AAV8-P301L tauopathy mice that change when near muscle features that make myofibers susceptible to circulating or propagating tau, such as blood vessels and nerve bundles, respectively. This finding suggests that tau has the potential to transcriptionally impact other organs beyond the central and peripheral nervous systems, and that mechanism may be mediated by proximity to circulatory or nervous systems^34^.

The transcriptional environment was different in myofibers of AAV8-P301L tau expressing mice. The transcript *Amd1* was significantly differentially expressed in 7 out of 8 region types analyzed. This metabolic polyamine biosynthesis enzyme and its pathway contribute to skeletal muscle force generation, although it is not clear to what extent ^35,36^. Transcriptional detection of spermine oxidase (*Smox*), also involved in polyamine biosynthesis, was differentially expressed in AAV8-P301L male myofibers proximal to nerve bundles, suggesting that the polyamine biosynthesis pathway may be altered. The consistent differential expression of *Amd1* makes the gene attractive as a biomarker. Furthermore, genes encoding proteins essential for generating force such as *Ache*, *Myh4*, and *Casq2* were altered. The upregulation of *Casq2* (1.18 Log2Fold Change) in AAV8-P301L males suggests compensation ensuring proper calcium buffering (as observed in the force frequency curves from the soleus and EDL). It is unclear if the transcriptional changes represent adaptive or pathologic responses, but the observed transcriptional changes suggest that skeletal muscle force production could be impacted.

*In vivo* weakness, mononuclear cell infiltration displacing contractile material, and transcriptional alterations alluded to the possibility that the muscle contractile machinery in AAV8-P301L tau mice was impacted. A reasonable explanation for muscle weakness is that tauopathy affects muscle innervation, thereby influencing force production. To independently study alterations in the contractile material of the muscle, we report specific force. Since the muscle is externally stimulated *ex vivo*, deficits in this measure are independent of motor neuron signaling and due to shortfalls in the contractile material of the muscle fibers generating force. In doing so, we discovered reduced specific force in the soleus muscle of AAV8-P301L tau males. This finding supports that tauopathy has peripheral consequences beyond gene expression that have functional detriments to skeletal muscle. We did not observe a deficit in specific force of the EDL. This could be explained by the normal heterogeneity in skeletal muscle. The soleus is a mixed fiber type muscle located in the posterior of the lower hindlimb and regularly recruited for posture and movement. The EDL is primarily a fast fiber type muscle located in the anterior region of the hindlimb, does not contribute to weight-bearing activity, and is infrequently recruited. Therefore, individual muscles may be impacted by tauopathy in a distinct manner and may be important to consider in clinical cases. This finding may be underpinned by differences in muscle usage since walking requires increased soleus functional demand relative to EDL function^37–39^. AAV8-P301L tau mice have increased activity detected by IR cage sensors, and the soleus may be susceptible to chronic over-usage if the increase in IR beam break activity correlates to walking and muscle usage. Our findings may also reflect the differences in susceptibility to aging phenotypes between the two muscles ^40^. The mice utilized in this study were collected at 15 months, and are not considered aged, but the tauopathy may induce aging phenotypes.

Beyond executing movement, skeletal muscle has essential roles in whole-body metabolism^41–43^. In tauopathy, it is unknown whether these functions are preserved in skeletal muscle. We observed consistent male-specific decreases in creatine kinase expression, indicating that high energy phosphate metabolism could be impacted. This supports claims that muscle metabolism is altered in tauopathy. Additionally, skeletal muscle is a primary site for glucose uptake/storage/release, and therefore significantly contributes to whole body metabolism. We observed a significant increase in *Ldha* expression in AAV8-P301L females, encoding lactate dehydrogenase A, a subunit of lactate dehydrogenase in skeletal muscle. This alludes to the preferential shift to anaerobic glycolysis and altered glucose utilization, or to the preferential production of lactate. Lactate transport has been postulated to exist from muscle fibers into circulation, and to neurons. This is enabled by monocarboxylate transporters that facilitate blood-brain barrier transport to allow lactate mediated oxidative metabolism as energy supply for the brain ^44–47^. This may be a meaningful consideration when there is high metabolic stress on neurons due to NFT accumulation. Brain metabolic stress mechanisms are active in tauopathies, such insulin signaling, inflammation, and ER stress ^8,9,48–50^. These stressors may call for increased metabolic demand on skeletal muscle to preserve homeostasis.

### Conclusion

Pathological tau species such as NFTs likely have far-reaching consequences in addition to neurodegeneration; our findings emphasize this in skeletal muscle. Tauopathy induces peripheral symptoms in the AAV8-P301L model and has the potential to influence additional skeletal muscle functions beyond generating force. Transcriptomic analyses indicate the potential for muscle-based biomarkers in assessing tauopathy risk, therefore positioning skeletal muscle as a source for tauopathy biomarkers in addition to blood-based biomarkers and imaging. Mechanisms underpinning reductions in force or mediating tau pathogenesis in distal muscles remain elusive and require additional efforts to understand how pathological tau mediates skeletal muscle anomalies, and the additional consequences imposed on skeletal muscle.

**Table 1.** DEGs in AAV8-P301L tau expressing mice.

## Acknowledgments

We thank the Abisambra and Esser laboratories for the helpful discussions and training. We also thank Dr. Todd Golde and Danny Ryu for their contribution of AAV constructs. This work was supported by NIH/NIA U01AG055137-S1 (K.E.), NIH/NIAMS R01AR079220 (K.E.), NIH/NIA R01AG074584-02 (J.A.), NIH/NIA R01AG075900-01 (J.A.), and Alzheimer’s Association AARG-D-21-847204 (J.A.).

## Author contributions

Conceptualization, B.A., K.E. and J.A.; methodology, B.A., K.E., and J.A.; investigation, B.A., G.H., and S.S.; writing – original draft, B.A.; writing – review & editing, B.A., G.H., K.E., and J.A.; funding acquisition, K.E. and J.A.; resources, S.P, K.E., and J.A.; supervision, K.E. and J.A.

## Declaration of interests

The authors declare no competing interests.

## Methods

### Mice

All animal studies were approved by the University of Florida Institutional Animal Care and Use and Committee and recombinant DNA use was approved by University of Florida EH&S Office. C57BL/6J mice were used in this study and did not have any notable physical abnormalities. Mice were maintained on food and water ad libitum and on a 12-h light/dark cycle. All mice were anesthetized and euthanized following IACUC-approved protocol at 15 months, and no physical abnormalities were detected. Brains were immediately collected and cut at the midsagittal plane, with one hemisphere fixed in 10% formalin and the other dissected to isolate cortical and hippocampal regions before being flash frozen in liquid nitrogen. Concurrently, the tibialis anterior, gastrocnemius, and soleus were collected and flash frozen. The contralateral tibialis anterior muscle was embedded in O.C.T. compound (Tissue-Tek 4583, Sakura Finetek USA, Inc.) and frozen in liquid nitrogen-cooled isopentane for cryo-sectioning. A cohort of AAV8-P301L tau expressing mice (N=7 males) and control mice (N=3 males) were anesthetized and euthanized at the Physiological Assessment Core at the University of Florida, in which under anesthesia the extensor digitorum longus (EDL) and soleus muscles were carefully dissected out for mechanical testing prior to collecting the brain or other muscles.

### AAV Injections

Recombinant AAV was produced and injected bilaterally into the ventricles of all neonatal mice (PND 0) as previously described ^51^. The construct expressing P301L-tau under the control of the human synapsin promoter was packaged in capsid serotype 8 for this study. Control mice were injected with either an empty AAV vector packaged in serotype 8, or with a construct expressing yellow fluorescent protein under the control of the human synapsin promoter packaged in serotype 8. All animals were bilaterally injected with 2 μL (1 x 10^13^ viral genomes) of AAV8 in each hemisphere.

### Grip strength and activity measures

Body weight of each mouse was measured and recorded at the time of the grip strength test. Forelimb grip strength was measured at 3 months of age following the TREAT-NMD SOP: DMD_M.2.2.001. Forelimb grip strength was measured 5 times with one minute rest periods in between. The average of the measures was used as the raw grip strength. Raw grip strength was then divided by body weight to calculate the grip strength normalized to body weight. Voluntary locomotor activity in the cage was continuously monitored using passive infrared motion detectors (Aurora; Ademco®) and the ClockLab analysis software (Actimetrics) in one-minute bins over 10 days. Locomotor activity data collected during the first two days were excluded to limit the detection of non-representative changes in activity from acclimation to a new cage.

### Muscle function measures

Under anesthesia, the EDL and soleus muscles were carefully dissected out at the Physiological Assessment Core at the University of Florida Wellstone/Myology Institute, and mechanical measurements (isometric tetanic force, force/frequency relationship, muscle weight) were performed following the TREAT-NMD SOP: DMD_M.1.2.002, as described previously (AU - Moorwood et al., 2013).

### Immunohistochemistry and histology

15-month-old hemibrains were fixed in 10% formalin overnight in 4°C upon dissection. Hemibrains were embedded in paraffin and serial sections (10μm) were cut from the midline.

Immunohistochemistry (IHC) was performed as previously described (Sakthivel et al., 2023), using the HT7 primary antibody (Thermo Fisher Scientific MN1000; 1:1000) and the corresponding Goat Anti-Mouse IgG (H+L) biotinylated secondary antibody (Vector Laboratories; 1:1000) to detect injected human derived tau with DAB labeling. The slides were imaged with the Aperio slide scanner (20x objective lens, Scan Scope^TM^ XT, Aperio Technologies, Inc. Vista, CA).

Tibialis anterior muscles embedded in OCT compound were cryo-sectioned into 8 μm thick cross sections. General myofiber morphology was visualized by staining the sections with hematoxylin and eosin and then imaged with the Aperio slide scanner (20x or 40x objective lens, Scan Scope^TM^ XT, Aperio Technologies, Inc. Vista, CA). After myofiber morphology was visualized, corresponding serial sections were chosen for additional IHC to demarcate myofiber perimeter. Cryo-sections were incubated for 1 hour in blocking buffer (5% Bovine Serum Albumin; 10% Normal Goat Serum; 1% Glycine; 0.1% Triton X-100; in TBS) and then labeled with anti-laminin antibody (Millipore Sigma L9393, 1:100) to demarcate myofiber perimeter diluted in blocking buffer and incubated overnight at 4°C. Following overnight incubation, sections were washed three times for 10 min each in blocking buffer. Corresponding secondary antibodies (ThermoFisher A-11012, Alexa Fluor^TM^ 594) were diluted in blocking buffer for 2 hours (1:500) and then washed 3 times for 10 min in blocking buffer. Immunofluorescent images were acquired and stitched automatically with the Keyence BZ-X710. Images with laminin stains were automatically processed using MyoVision 2.0 software to acquire fiber cross sectional area and minimum feret diameter as previously described ^54^.

### Biochemical fractionation of mouse brains and western blotting

Cortical brain regions isolated from a hemibrain, that were quickly frozen in liquid nitrogen and stored at -80°C were used for biochemical analysis. Each cortical sample was weighed and homogenized in ten volumes (10x the weight) of homogenizing buffer (50mM tris base, 10% glycerol, 2% SDS, 2% 2-mercaptoethanol, pH 8.8, protease and phosphatase inhibitors) with a Dounce homogenizer. The homogenates were fractioned to obtain a supernatant fraction with soluble tau (S1) and a pellet with sarkosyl-insoluble tau (P3) as previously described ^55^. Sarkosyl-insoluble pellets (P3) were resuspended in 50 μL of 4 x Laemmli sample buffer (Bio-Rad), separated into 10 μL aliquots, and stored at -80°C until used for western blot analysis.

Soleus tissue lysates were obtained by homogenizing the tissue in 50 volumes (50x the weight) of homogenizing buffer with a Dounce homogenizer. Samples were heated at 80°C for 5 min and were centrifuged for 15 min at 20,000g to remove cellular debris. The supernatant was collected and stored at -80°C until used for western blot analysis. A Bicinchoninic acid protein assay was performed on the S1 brain samples and soleus supernatant. 10 ug of protein from cortical samples and 45ug of muscle samples were diluted in 4x Laemmli sample buffer (Bio-Rad). P3 brain samples were already directly resuspended in 4x Laemmli sample buffer. All samples were heat-denatured for 5 min at 100°C and run on a 4-20% Criterion Tris-HCl Protein gel and transferred to PVDF membrane (Millipore). Membranes with muscle tissue homogenates were blocked in 5% non-fat dry milk in TBS/0.2% Triton X-100 and incubated overnight in Tau12 (Millipore Sigma; 1:1000) and actin primary antibody (Millipore Sigma; 1:1000) diluted in 5% milk in TBS/0.1% Triton X-100 rocking at 4°C. Membranes were incubated in Goat-Anti-Mouse HRP-conjugated secondary antibody (SouthernBiotech; 1:2000) for 1 h at room temperature and detected by ECL (PerkinElmer). Membranes with cortical samples were blocked in Intercept (TBS) blocking buffer (LI-COR) rocking overnight at 4°C, and incubated for 1h in either Tau12 (Millipore Sigma MAB2241; 1:1000), Tau46 (Santa Cruz Biotechnology Inc. sc-32274; 1:1000), or AT8 primary antibody (Thermo Fisher Scientific MN1020; 1:1000) diluted in TBS/0.2% Tween-20. Membranes were incubated with the corresponding Goat Anti-Mouse IgG (H+L) IRDye secondary antibody (LI-COR 926-32212; 1:12,500) for 1 h at room temperature and imaged on an Odyssey M Imager (LI-COR).

### RNA Isolation and NanoString Mouse Whole Transcriptome Atlas (WTA) assay

In order to determine the RNA integrity number (RIN) for each sample, RNA was isolated from flash frozen 8 μm thick sections of tibialis anterior muscles embedded in OCT using the RNeasy Mini Kit (Qiagen) and evaluated using the RNA high sensitivity kit (Agilent RNA 6000 Pico Kit) and an Agilent 2100 Bioanalyzer. Only samples with a RIN >7.6 were chosen for the Nanostring Mouse Whole Transcriptome Atlas (WTA). The mouse WTA assay (NanoString Technologies) was performed as previously described ^56^ following the manufacturers protocol and NanoString University supplemental protocols with slides containing 8 μm thick OCT cryosections of the tibialis anterior muscle. A primary-conjugated anti-laminin antibody (Novus, 1:500) and SYTO13 (NanoString Nuclear Stain Morphology kit) were used as morphology markers. For each of the 4 groups (tauopathy male or female and control male or female), 3 biological replicates of the cross sections were used. From each cross section (12 cross sections total), regions were chosen. 4 region types were chosen (standard fibers, regions with fibers containing central nuclei, regions of fibers proximal to nerve bundles and regions of fibers proximal to vasculature) on each cross section. 1-2 replicates of each region type was chosen by the user, depending on the frequency of certain muscle hallmarks in the section (regions with central nuclei, blood vessels, or nerve bundles, all had minimum one region). Every selected region was ∼9000 μm^2^ in area. After collection of ROIs, libraries were prepared as described in protocol and submitted for sequencing to the UF Interdisciplinary Center for Biotechnology Research. FASTQ files were then converted into DCC files using BaseSpace and analyzed using the Nanostring Data Analysis suite.

### Statistical analysis

GraphPad Prism 9 software was used to perform statistical tests. Results are reported as means ± standard error of the means. Unpaired Student’s t-tests were performed for comparisons between AAV8-P301L tau expressing mice and controls of the same sex and of the same parameter. A two-way ANOVA was used to make multiple comparisons between tauopathy mice and controls of both sexes. *P < 0.05* was considered significant.

Data from the NanoString Mouse Whole Transcriptome Atlas assay was analyzed on the GeoMx DSP analysis suite (NanoString). A linear mixed model with Benjamini-Hochberg correction was performed to acquire unadjusted and adjusted p-values. Targets were considered of interest with either an adjusted P value < 0.1, or an unadjusted P-value <0.05 and a Log2Fold Change of +/-1.

## Lead contact

Additional information and requests for resources should be directed to and will be fulfilled by the lead contact, Jose Abisambra.

## Materials availability

This study did not generate novel reagents.

## Data and code availability

All data reported in this paper, or additional information required to reanalyze the data reported will be shared by the lead contact upon request.

This paper does not report original code.

## Notes

### Competing Interest Statement

The authors have declared no competing interest.

