## Supplementary figures and images for "Targeted brain-specific tauopathy compromises peripheral skeletal muscle integrity and function"

### Fig. S1

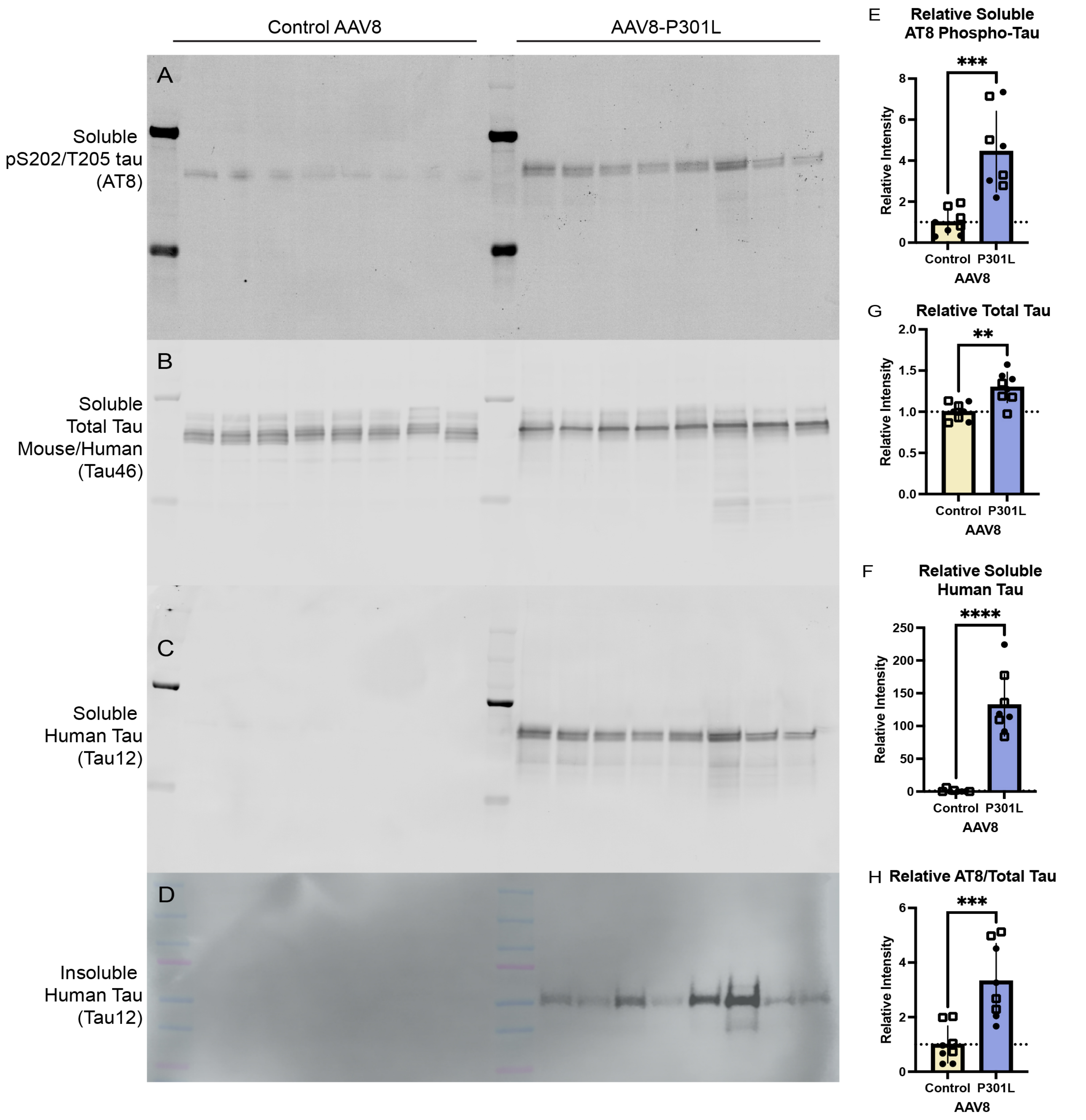

### Fig. S2

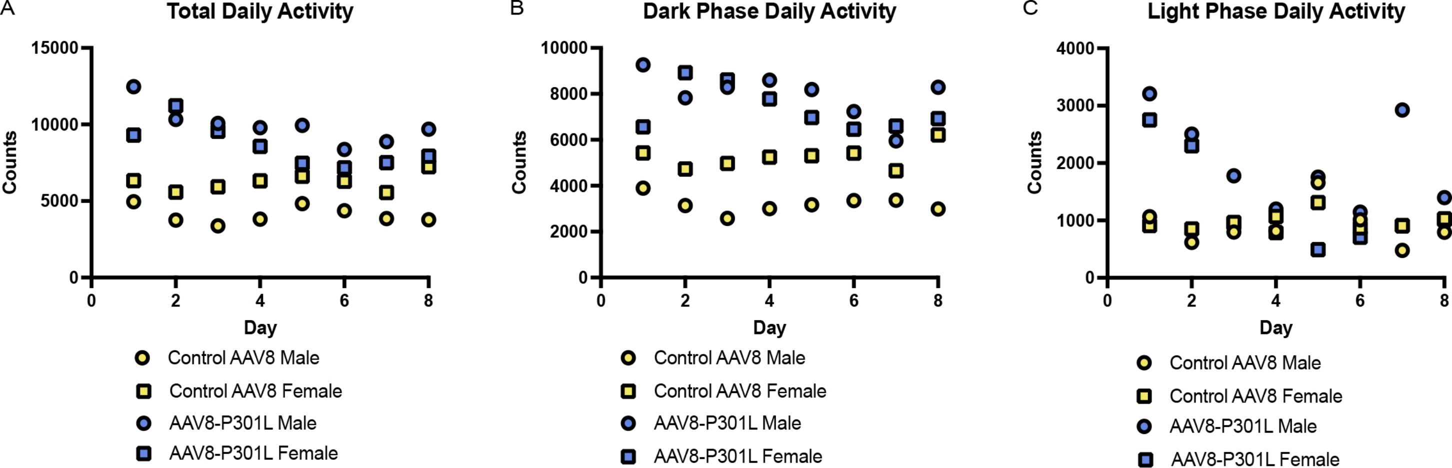

### Fig. S3

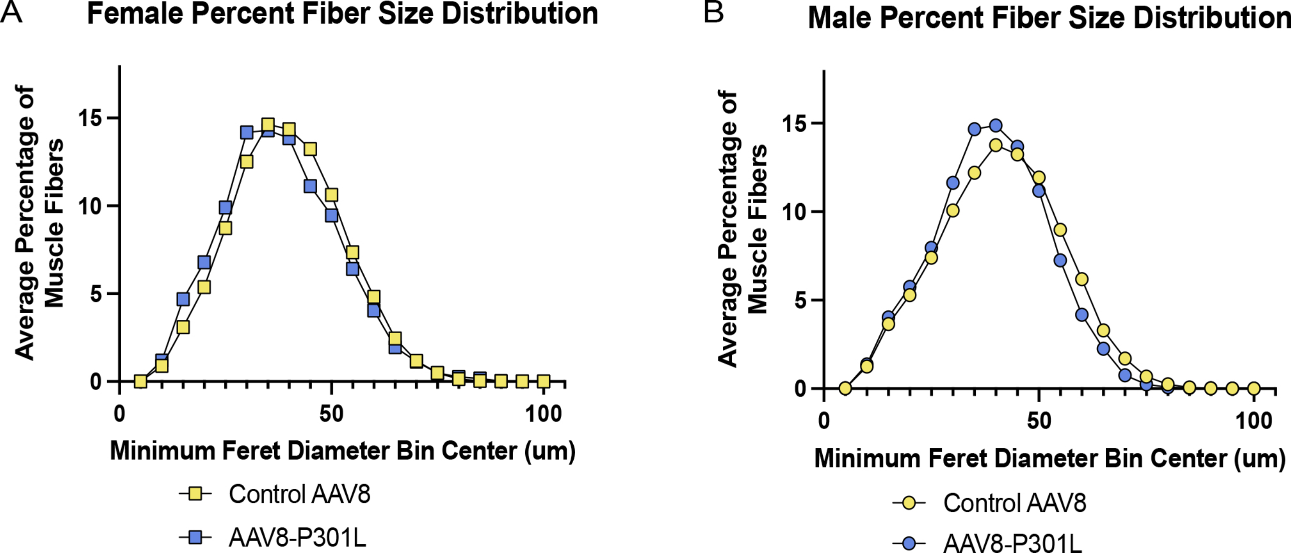

### Fig. S4

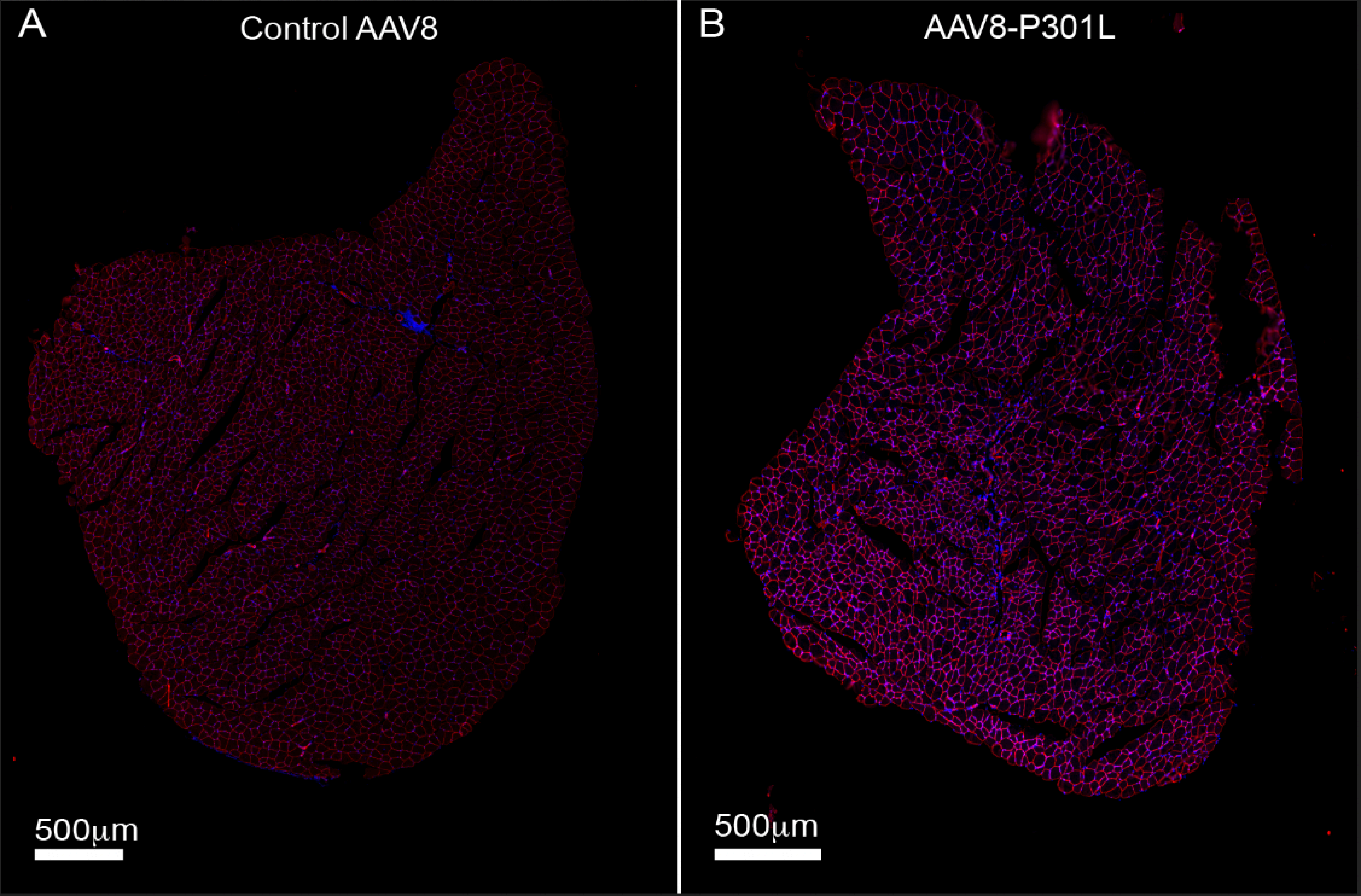
